# Knee Biomechanics during Walking in Individuals with Anterior Cruciate Ligament Repair: The Role of a Custom 3D Printed Knee Brace

**DOI:** 10.1101/2024.07.04.601890

**Authors:** Florian Mougin, Mickaël Begon, Gauthier Desmyttere, Jacinte Bleau, Marie-Lyne Nault, Yosra Cherni

## Abstract

**Background:** Anterior cruciate ligament (ACL) injuries frequently lead to altered gait biomechanics and muscle activation patterns, increasing the risk of osteoarthritis. Knee braces are commonly used to address these issues although a lack of consensus remains regarding their clinical benefits. The recent emergence of 3D-printed braces, lighter and personalized, could improve rehabilitation.

**Objectives:** To evaluate the effect of a novel custom-made 3D-printed knee brace (Provoke™) in individuals after unilateral ACL reconstruction during walking. The brace incorporates an asymmetrical hinge system aimed at stabilizing the knee joint while minimizing compensatory movements.

**Methods:** Fourteen participants with unilateral ACL reconstruction wore the Provoke™ brace while walking at comfortable and fast paces. Knee kinematics and kinetics, and muscular activity (rectus femoris, vastus medialis, and semitendinosus) were assessed with and without the brace. Two-tailed non-parametric paired T-tests were used to assess the biomechanical effect of the brace.

**Results and conclusions:** The Provoke™ brace improved knee kinematics, facilitating a more neutral knee position by reducing valgus angles (-1.95°), and increasing flexion angles (+1.14°). Additionally, it enhanced muscle activation, particularly of the rectus femoris, suggesting improved quadriceps function. Overall, the Provoke™ brace effectively improves knee function and reduces muscle imbalances in individuals undergoing ACL reconstruction. It may help prevent further injury and reduce the risk of post-traumatic osteoarthritis development. The long-term effects of brace use in ACL rehabilitation must be investigated.

**Key Points:**

- Walking while wearing Provoke™ brace allow to reduce knee valgus angles which could counteract the development of post-traumatic osteoarthritis.
- Braced walking may mitigate the stiffened knee gait strategy by increasing the peak knee flexion angle during the stance phase of the gait.
- Increased rectus femoris activation during early stance phase could improve knee function and stability, acting against quadriceps weakness often observed after anterior cruciate ligament reconstruction.

## INTRODUCTION

Anterior cruciate ligament (ACL) rupture is one of the most common knee injuries which prevails among young individuals and elite athletes ^1^, causing significant challenges to patient’s health and function. Approximately 200,000 ACL injuries are reported annually in the United States, and 2 million worldwide, making it a prior subject for research and investment ^2,3^. ACL rupture leads to knee instability, particularly in the sagittal plane, and a loss of knee joint proprioception which affects control mechanisms and may impact everyday tasks such as walking ^4^. Treatment ranges from conservative therapy to reconstructive surgery which stands as the prevailing approach when managing ACL injuries ^5^.

While ACL reconstruction (ACLR) surgery offers a pathway to functional recovery, it also carries risks. Pathological compensation movements after ACLR often alter gait in both legs^6^. To avoid instability and discomfort, patients might adopt a “stiffened leg” walking strategy, reducing knee range of motion in flexion-extension ^7^ and leading to imbalances in muscle recruitment. A decreased activation of the extensor muscles (quadriceps avoidance) has also been reported ^8,9^. Furthermore, it has been reported that six months after ACLR, individuals may develop adaptative dynamic knee valgus (40% of lower limb malalignment compared to baseline)^6^. However, this adaptation is a common cause of increased joint rate of health degeneration leading to post-traumatic osteoarthritis ^2,7,10^. Moreover, it not only increases by sixfold the risks of ACL re-injury within the first year or upon return to sports, but also the risk of contralateral injury ^6^. To mitigate these issues, patients frequently undergo rehabilitation protocols after ACLR, including knee bracing to safeguard the graft ^11^, support joint stabilization, increase confidence ^2,12,13^, and reduce the risk of reinjury, especially for skiers ^4^.

Knee braces may improve sagittal and frontal plane biomechanics by **1)** increasing the knee flexion angle and thereby minimizing the effect of the stiffened knee gait strategy ^7^ and **2)** limiting aberrant frontal plane motion partly responsible for an increased risk of over-injury ^6^. However, brace efficacy remains controversial due to the lack of consensus on their potential clinical benefits. In their meta-analysis, Yang et al. ^13^ did not report any established advantage regarding the application of braces in knee function, pain, and stability assessments when compared to an unbraced condition. Furthermore, post-surgery knee bracing might be linked with some drawbacks such as increased thigh muscle atrophy compared to individuals without braces, potentially compromising long-term muscular joint stabilization, proprioception, and neuromuscular control ^2,13^. Wearing conventional knee braces can also result in discomfort, which may enhance gait deviations over time. Custom-made braces are superior in terms of fit, pain reduction, and response to disability ^14^. However, their manual fabrication by qualified orthotists is expensive and time-consuming ^14,15^. Additive manufacturing technology has recently revolutionized the creation of personalized braces, offering quicker and more cost-effective solutions tailored to the patient’s specific anatomical and pathological features. Few studies have evaluated the benefits of 3D-printed knee braces and the literature lacks evaluation of their biomechanical effectiveness, relying predominantly on subjective criteria such as wearing comfort and pain assessment ^13,14^.

The present study aimed to evaluate the effects of a new generation custom-made 3D-printed knee brace on the knee biomechanics and muscular activity in individuals with a unilateral ACLR during walking. Three hypotheses were formulated: 1) the custom-made 3D-printed knee brace induces an increase in knee flexion during stance phase and enhance knee stabilization in the frontal plane by minimizing excessive varus-valgus angles; 2) Quadriceps electromyography (EMG) activity increases when wearing the brace counteracting “stiffened” gait strategy; and 3) Fast-paced walking amplifies the effect of the brace on injured leg kinetic, kinematic and EMG.

## METHODS

### Participants

The inclusion criteria were: (1) 20 to 60 years old, (2) have undergone a unilateral ACL reconstruction (3) use a knee brace routinely, (4) no lower limb surgery or injury during the last three months, and (5) able to walk at least 3 minutes with or without brace. The protocol was approved by The Research Ethics Board of Université de Montréal. All participants gave their written informed consent before data collection.

### Experimental protocol

The Provoke™ brace (Osskin Inc, Montreal, Canada) was used for this study. It consists of a 3D-printed custom knee brace designed for physically active individuals and athletes to address various ligament conditions including ACL injury. This brace features a unique asymmetrical hinge system that aims to mimic the knee’s “screw-home” natural motion, intending to stabilize the joint while minimizing compensatory movements and preventing extreme internal-external rotation that can be seen in ACLR patients ^16^. In healthy knees, the ACL is the guide for this screw-home mechanism ^16^. To mitigate frontal plane issues, the brace is engineered to diminish abnormal valgus angles and redistribute valgus forces to the strengthened section of the brace. This 3D-printed brace (270g) is lighter than conventional braces ^14^ and aims to provide a precise fit, comfort, and effective transfer of valgus loads. Participants had at least three weeks of accommodation before data collection.

During the evaluation session, participants started with a 3-min walking familiarization on the instrumented treadmill (Bertec, Columbus OH, United States). Participants self-selected comfortable and fast paces were determined during this phase. Then, participants walked at their self-selected comfortable and fast paces for 30 seconds under two conditions: with and without brace in a randomized order.

### Data recording and processing

#### Kinematic and Kinetics outcomes

An 18-camera motion capture system (T40S VICON, Oxford, UK) measured the 3D trajectories of reflective markers placed on participants’ lower limbs (Figure 1) while ground reaction forces were measured under each foot using a dual belt instrumented treadmill (Bertec, Columbus OH, United States). A seven-segment (pelvis, 2x thigh, 2x shank, 2x foot), 42 degrees-of-freedom model was used to determine the joint kinematics. Joint centres and axes were calculated based on functional movement trials performed pre-testing for each participant using the SCoRE algorithm ^17^. Joint angles were calculated using inverse kinematics ^18^ with the following angle sequence: flexion-extension, abduction-adduction, and internal-external rotation. The zero angular positions were defined using a static trial in anatomical position. Net joint moments were calculated using inverse dynamics, based on joint kinematics, ground reaction forces at the foot, and the inertial properties from De Leva, P ^19^. Thus, joint moments were expressed in the same coordinate system as the kinematics. Kinematic data were collected at 100 Hz and ground reaction forces at 2000 Hz. For joint angles and moments, 15 continuous gait cycles of the injured leg were analyzed under each condition. A 5% body weight vertical cut-off force threshold was applied to distinct steps on force plates. Gait cycles were normalized on a 0-100% scale beginning and ending at heel strike. All kinematic and force data were filtered using a zero-lag 2nd order Butterworth low-pass filter, with a cut-off frequency of 10 Hz. Joint moments were normalized to the participant’s body mass. During the analysis of fast walking, one participant was excluded from the kinematic data and two from the kinetic data due to reconstruction issues.

**Figure 1:**
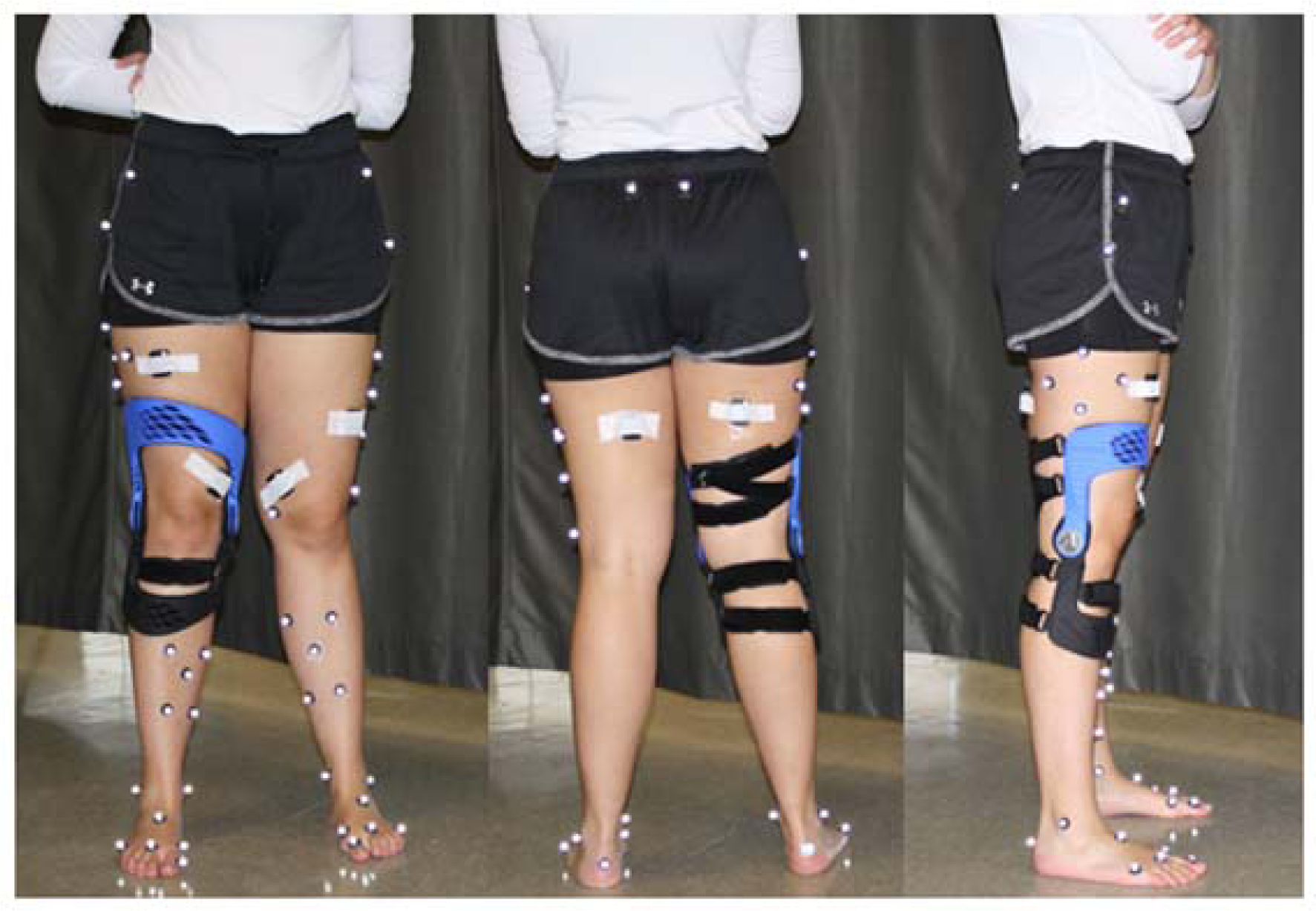
Placement of Markers and EMG sensors.

#### Electromyography

Three EMG electrodes (Delsys Trigno Wireless system) were positioned on the rectus femoris (RF), vastus medialis (VM), and semitendinosus (ST) muscles (Figure 1). The recommendations of Surface EMG for Non-Invasive Assessment of Muscles (SENIAM) were followed for suitable skin preparation and electrode placement ^20^. EMG data were sampled at 2000 Hz. Raw EMG data were 10-400 Hz band-pass filtered, full-wave rectified, and low-pass filtered at 9 Hz to obtain the linear envelope. All filters were 4th order, zero lag Butterworth filters. EMG data were time normalized (0-100%) according to the gait cycle. Then, the average muscle activation profile for each subject was estimated by averaging the linear envelope of all individual gait cycles. Finally, the EMG signals of each muscle, for each patient, were normalized to the peak signal of the contralateral limb.

### Statistical analysis

Descriptive results are expressed as mean (±95% Confidence interval (CI)). Paired *t*-tests were used to evaluate the knee brace effects on participants walking pace. Two-tailed non-parametric paired T-tests using Statistical non-Parametric Mapping (SnPM) ^21^ were used to assess the knee brace effect on the time histories of joint angles, joint moments, and EMG envelopes (over the entire gait cycle) during both walking conditions. The *t*-test was performed over the entire gait cycle to form intervals over which significant differences existed. Bonferroni correction was applied to the kinematic, kinetic, and EMG data resulting in a corrected p value of 0.002. Cohen *d* was calculated to obtain the effect size of each significant difference between the two conditions. Effect sizes in the range of 0.0-0.3, 0.3-0.7, 0.7-1.0 as well as their negative peers were considered as weak, moderate, and strong effects, respectively ^22^.

## RESULTS

Fourteen participants (age: 48.1 ± 10.9 years; weight: 70.4 ± 11.2 kg; height: 174.9 ± 6.6 m; sex = 4 Females) were included in the study. Wearing the brace did not significantly affect the comfortable walking pace (0.91±0.18 vs 0.91±0.17 m/s without brace; p=0.97) nor the faster pace (1.29±0.16 m/s vs 1.25±0.17 m/s without brace; p=0.09).

### Knee kinematics and kinetics

Wearing the brace had a significant effect on knee kinematics at both walking paces (Figure 2 and 3). When the brace was worn, participants’ <u>knee flexion</u> increased throughout the stance phase (p=0.001), with a weak effect size, corresponding to an average increase of 1.14° and 1.40° for normal and fast walking. Those kinematic changes were accompanied by a weak increased knee flexion moment after heel strike for both walking paces (0.15<d<0.28).

**Figure 2.**
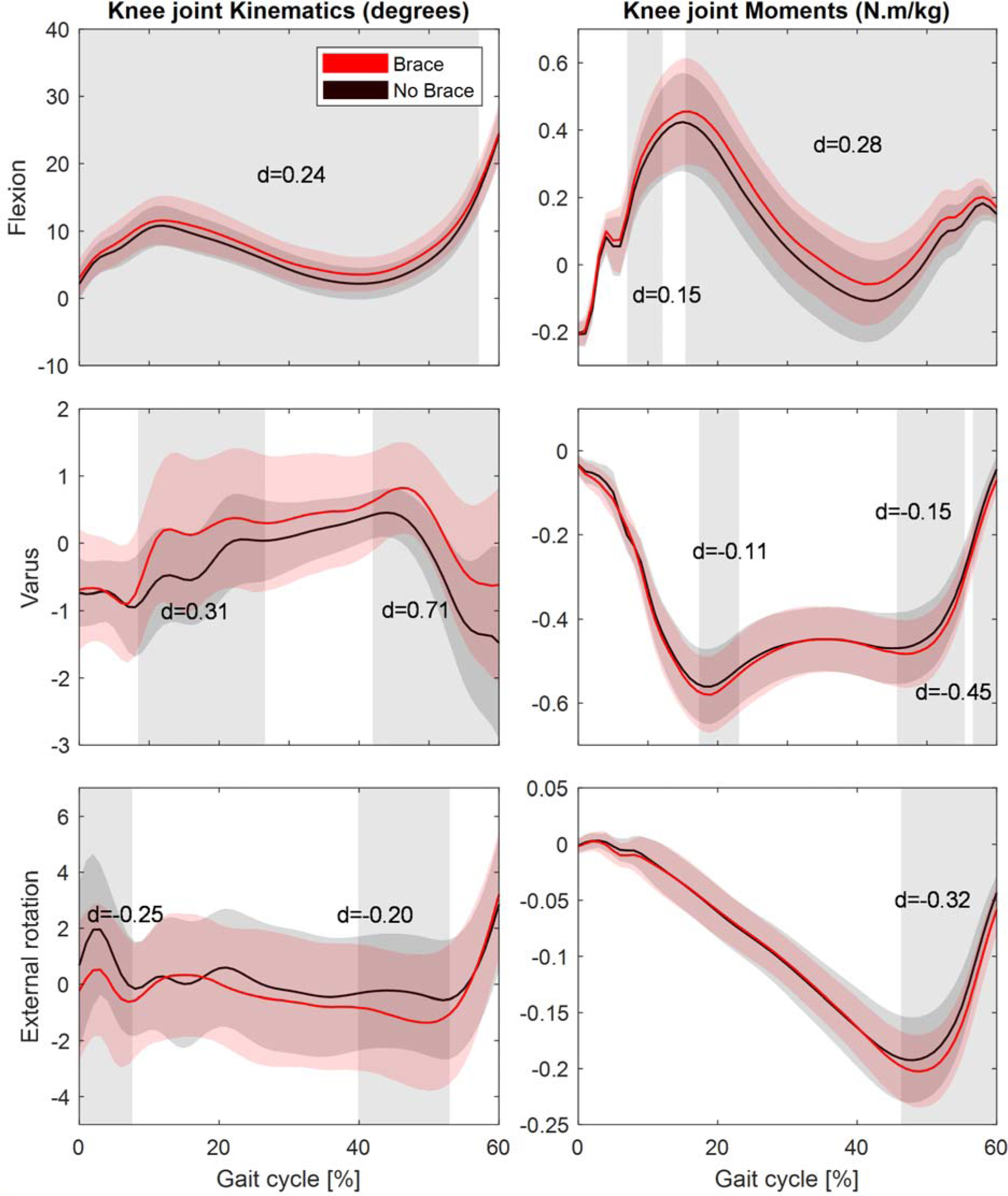
Knee joint angles and moments during the stance phase (0-60%GC) on the ACLR leg during normal walking. Brace condition is shown in red, and no brace in black. The shading represents the standard deviations. The grey bands represent periods of significant difference between the conditions with their mean Cohen d.

**Figure 3.**
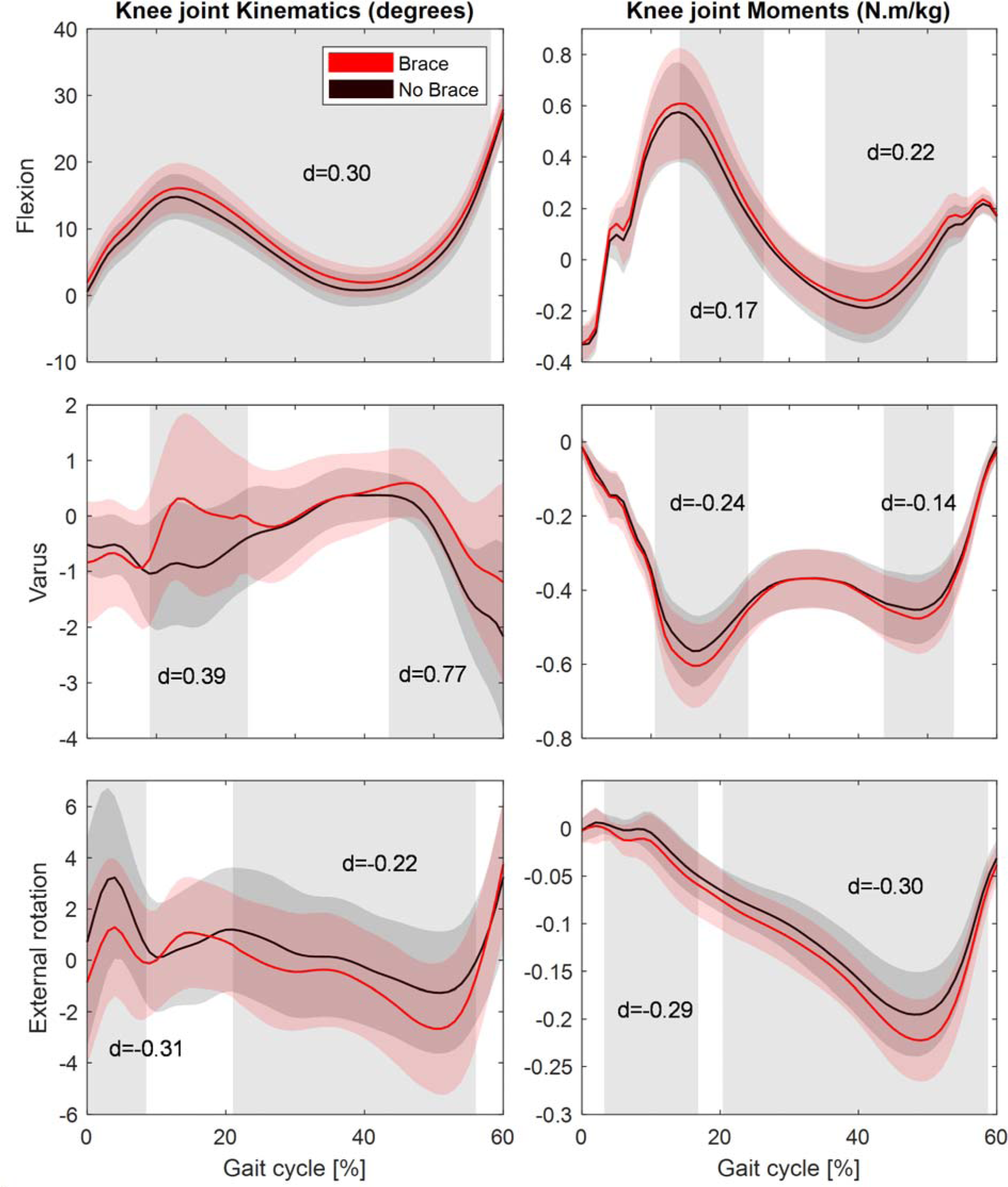
Knee joint angles and moments during the stance phase (0-60%GC) on the ACLR leg during fast walking. Brace condition is shown in red, and no brace in black. The shading represents the standard deviations. The grey bands represent periods of significant difference between the conditions with their mean Cohen d.

The <u>valgus angle</u> was significantly decreased in the brace condition for both walking paces (p=0.001), especially during the loading phase with moderate effect size (p=0.001; 0.31<d<0.39) and from pre-swing to swing phase with strong effect size (p=0.001; 0.71<d<0.77) with average decreases of 1.95° and 2.39°, respectively. Peak significant differences between no brace and brace conditions were seen at the swing phase with 4.83° and 5.36° for walking and fast walking respectively (p=0.001). The decrease in valgus angle was also associated with a weak to moderate increase in valgus moment for both walking paces (-0.45<d<-0.11).

A shift towards <u>internal rotation</u> was seen in both walking paces (p=0.001) when wearing the brace. The increase in internal rotation angles was weak to moderate for both walking and fast walking (-0.31<d<-0.20). These changes were associated with a weak to moderate increased internal rotation moment at both walking paces (-0.32<d<-0.29).

Regarding the contralateral limb (Appendices 1 and 2), no significant difference in knee kinetic and kinematic was observed when wearing the brace on the injured limb, except for a weak decrease in valgus moment for both walking pace (p=0.001, 0.10<d<0.19) with the brace. A weak increase in external rotation angle for walking (d=0.12), and a weak increase in flexion angle for fast walking (d=0.11).

### Muscle activity

During comfortable walking (Figure 4), a significant increase in rectus femoris activation was observed in the injured limb (p=0.001) with a weak to moderate effect size (0.23<d<0.36). Regarding the vastus medialis, a decrease was observed between 77−83%CG (p=0.001, d=-0.42). During brace condition, a decreased semitendinosus activity was observed between 40-51%GC with an effect size of d=-0.29.

**Figure 4.**
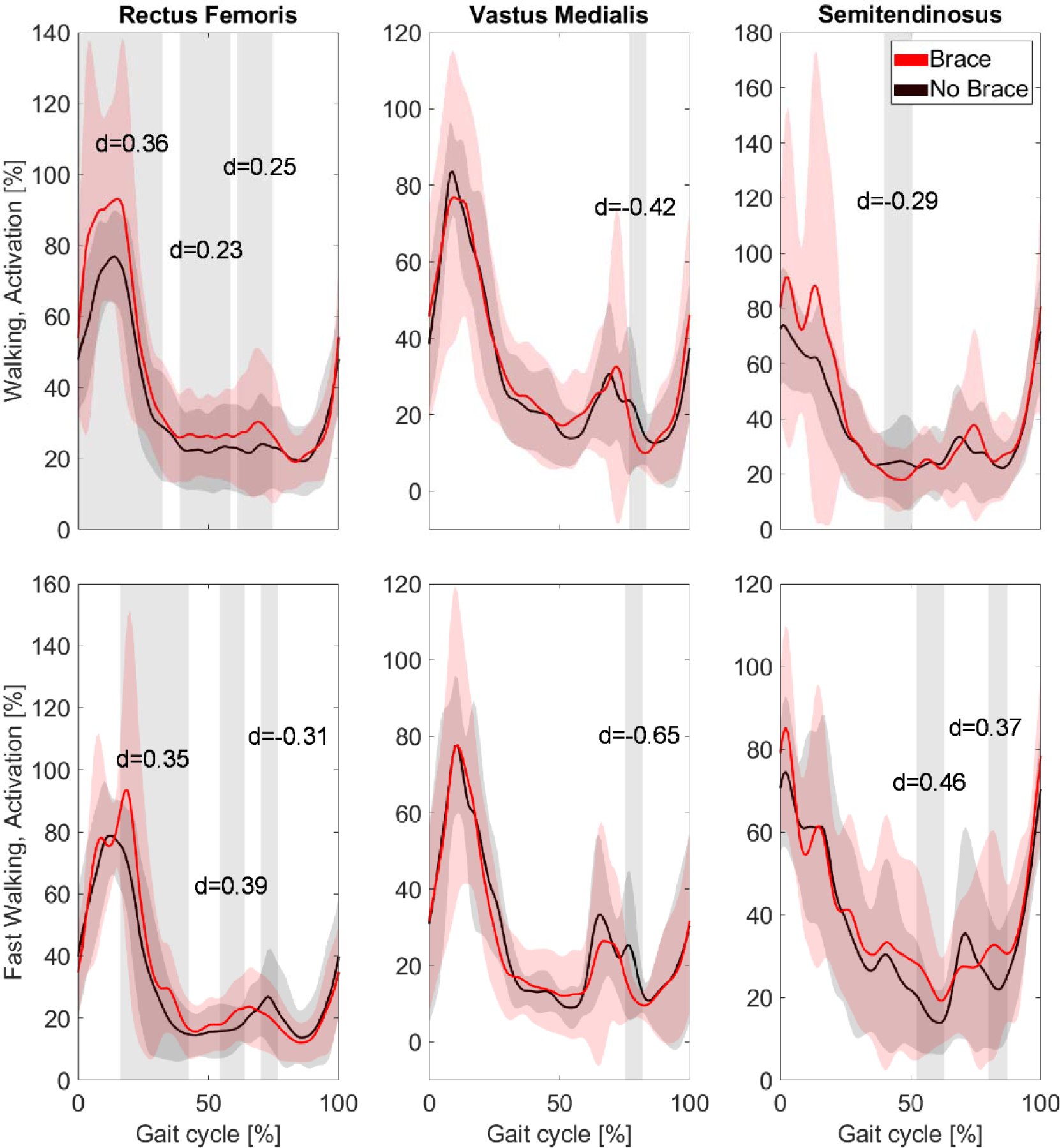
EMG activities during normal walking and Fast Walking tasks. Brace condition is shown in red, and no brace in black. The shading represents the standard deviations. The grey bands represent periods of significant difference between the conditions with their mean Cohen d. EMG is normalized with respect to peak value of the contralateral limb.

At <u>the fast-walking pace</u>, increases in rectus femoris activation (p=0.001) on the injured leg were observed associated with moderate effect size between 16-43% and 54-64%GC. A significant decrease in vastus medialis activity was observed with a moderate effect size of d=-0.65 between 75-82%GC. Semitendinosus activity increased significantly with moderate effect sizes (p=0.001, 0.37<d<0.46) between 52-63% and 80-87%GC. On the contralateral limb (Appendix 3), significant decreases in rectus femoris activation (p=0.001) were seen during the swing phase with moderate effect size for both walking paces. Weak to moderate decreases in vastus medialis activation (p=0.001) were observed during normal and fast walking (-0.68<d<-0.13). Significant increases in ST activity (p=0.001) at heel strike were observed during both walking paces, followed by a decrease at fast walking (all moderate effects).

## DISCUSSION

This study assessed the impact of a novel custom-made 3D-printed knee brace on knee biomechanics in individuals who have undergone unilateral ACLR, during walking. Consistently with our hypothesis, the results reveal that the Provoke^TM^ brace reduced knee valgus angle and slightly increased valgus moment, placing the knee in a more neutral position. Increased knee flexion angle and rectus femoris activity during the stance phase was also reported when wearing the brace.

### Knee kinematic and kinetic

The most probable mechanism of ACL re-injury appears to be valgus-rotational loading ^23^. Based on our results, the Provoke^TM^ brace facilitated a more neutral knee valgus angle during both walking paces (i.e., reduction >1.95°). The brace could reduce the risk of ACL re-tear by preventing valgus malalignment and extreme tibial rotation, both resulting in ACL strain ^24^. Additionally, valgus malalignment increases the risk of knee post-traumatic osteoarthritis early development and progression ^25,26^, with a 30% prevalence within 5-year ^10^. Felson et al. ^25^ reported that mild to moderate valgus malalignment, ranging from 1.1° to 3.0°, was associated with an increased risk of osteoarthritis progression. Our findings indicate a significant reduction in valgus (i.e., reduction >1.95°) comparable to that seen in Felson’s study, suggesting that the Provoke^TM^ brace could have the potential to attenuate the progression of post-traumatic osteoarthritis. In such cases, previous research has shown that wearing an unloader knee brace benefits cartilage health ^7,27^. Hart and colleagues^10^ observed increased peak knee flexion (+3.2°), varus (+1.7°), and decreased peak internal rotation (-3.0°) angles when participants wore a conventional (non-custom) unloader knee brace to correct valgus malalignment following ACL reconstruction ^10^. They also noted an increased peak knee flexion, adduction, and internal rotation moments during stance phase. Consistent with these previous studies, our findings suggest that the Provoke™ brace primarily functions as an unloader knee brace, targeting the frontal plane to reduce lateral knee loads and achieve a more neutral leg alignment to protect against joint deterioration. The shift towards internal rotation when wearing Provoke^TM^ could counteract the fact that abnormally increased tibial rotation, which persists after ACLR, is significantly associated with the development of new articular cartilage lesions at a mean of 8.4 years after reconstruction ^28^. Regarding muscle activity, in contrast to ^29^ study, we found a significant increase in rectus femoris activation with the Provoke^TM^ brace during stance and early swing phase at comfortable walking pace. Moreover, knowing that the brace is bespoke and extremely light, it could increase wearing adherence and enhance the benefits over time. Retrospective studies of ACL-injured patients have also revealed an association between increased valgus moments during gait and the development of knee osteoarthritis ^30,31^. However, walking with Provoke^TM^ brace did not induce a significant decrease in knee valgus moment. It could be deduced from the combination of a more neutral frontal plane posture, along with the almost consistent valgus moments, that the brace is absorbing some of the valgus loading, since moments calculated by inverse dynamics correspond to efforts sustained by the knee plus the brace. Nevertheless, the current study design limits our ability to precisely determine how loads are distributed among specific substructures or areas.

The observed increase in knee flexion (i.e., increase>1.14°) angle during the stance phase of walking with the brace may suggest improved knee joint function and mobility potentially because confidence get higher with the knee abduction and rotation stabilization. This effect might counter the stiffened knee gait strategy where the injured leg is kept close to the centerline, a strategy reported as detrimental to knee joint integrity ^2,7^. This could lead to enhanced gait mechanics and overall functional performance, ultimately aiding in the rehabilitation process for ACLR patients. This finding aligns with the conclusions of Evans-Pickett and colleagues^7^, who proposed that bracing may alleviate the stiffened knee gait strategy. Such an effect could potentially mitigate the onset of post-traumatic osteoarthritis once more.

### Muscle activities

Improved quadriceps activation and strength are essential for knee stability and dynamic movement control in ACL management ^4^. Our results reveal an average 8.5% increased RF activity during walking while wearing the brace. After ACL injury, the rationale behind the stiffening strategy is to limit or control undesirable anterior translation of the tibia during gait, a function typically performed by the intact ACL. However, identified imbalances in knee muscle strength predispose patients to re-rupture, underscoring the significance of restoring muscular strength for returning to pre-injury levels of sports activity ^4^. Quadriceps weakness, often seen after ACLR ^32^ (i.e., up to 30% decrease in quadriceps strength in the reconstructed limb when compared to the contralateral limb 6 months after surgery ^33^) is also suggested as an aggravating risk factor of knee osteoarthritis ^31^. In this context, Provoke^™^ brace use may hold promise in mitigating re-injury risks via increasing quadriceps recruitment and restoring the lower limb muscular balance. However, it is noteworthy that the activity of other key muscles such as the vastus medialis and semitendinosus remained relatively stable. The slight decreases in vastus medialis activation in the swing phase, could be considered suboptimal in the long run, given its role in maintaining proper patellar tracking. Indeed, prolonged knee bracing could result in muscle atrophy ^4,13^. In this context, findings from ^34^ suggested that the daily usage of a valgus unloader brace, averaging 4.7 hours per day reported by patients, had either no effect or even greater improvement on muscular strength during maximum voluntary isometric contraction against a dynamometer over a 6-month period compared to baseline. This suggests that prolonged brace use may not greatly contribute to vastus medialis muscle atrophy, especially considering its relative greater activation observed in the injured leg at specific points in the gait cycle under both brace conditions. Also, thanks to technological advancements, next-generation braces might challenge conventional beliefs about muscle atrophy. Clinical studies of the Guardian OA Rehabilitator™ Knee Brace show significant improvements in quadriceps and hamstring strength after wearing the brace for at least 3 hours a day for 90 days ^35^. For specific or challenging activities, the constated change in muscle recruitment caused by the Provoke™ could present real benefits for knee stability and proprioception. Moreover, its benefits could be enhanced by quadriceps strength training (i.e., enhance neuromuscular control, prevent re-injury, prevent muscle atrophy) which emerges as a cornerstone of ACLR rehabilitation ^4^.

Overall, the findings of the present study align with previous studies ^7,10,12^ highlighting the potential of knee braces in improving biomechanical parameters and, in this case, muscle activation patterns in ACLR patients. From a functional point of view, the Provoke^TM^ brace can certainly address some of the gait adaptations seen in people who have undergone ACLR, such as reduced knee flexion, valgus misalignment, and quadriceps avoidance. Also, new innovative functional knee brace designs could offer superior knee joint protection compared to conventional braces, aiding in post-injury or post-surgery rehabilitation, ultimately improving patient outcome while providing significant economic benefits ^35^.

### Study limitations

Some limitations should be acknowledged. Firstly, this study included a relatively small sample size and lacked a control group. While the chosen tasks represent low to moderate-intensity daily living movements, it is suggested that at-risk motion in ACLR patients primarily manifests during high-intensity activities ^16^. Consequently, our study’s task selection may not fully assess the brace’s effectiveness or limitations. Finally, it should be noted that our study did not gather information on the surgical techniques used for reconstruction among the individuals which affect the variability in gait alterations in ACLR individuals ^10^.

## CONCLUSION

Our study underscores the potential of the Provoke™ knee orthosis in improving knee joint biomechanics and muscular activity in ACLR patients. This brace may improve post-ACL rehabilitation by reducing the risks of reinjury and osteoarthritis development by addressing key compensations in ACL-reconstructed individuals. While further research is warranted to assess the long-term effects of using the Provoke™ brace in ACL rehabilitation, our results provide valuable insights showing the potential of 3D printed custom brace in optimizing rehabilitation strategies and improving patient outcomes.

## Supplemantary materials

**Appendix 1:**
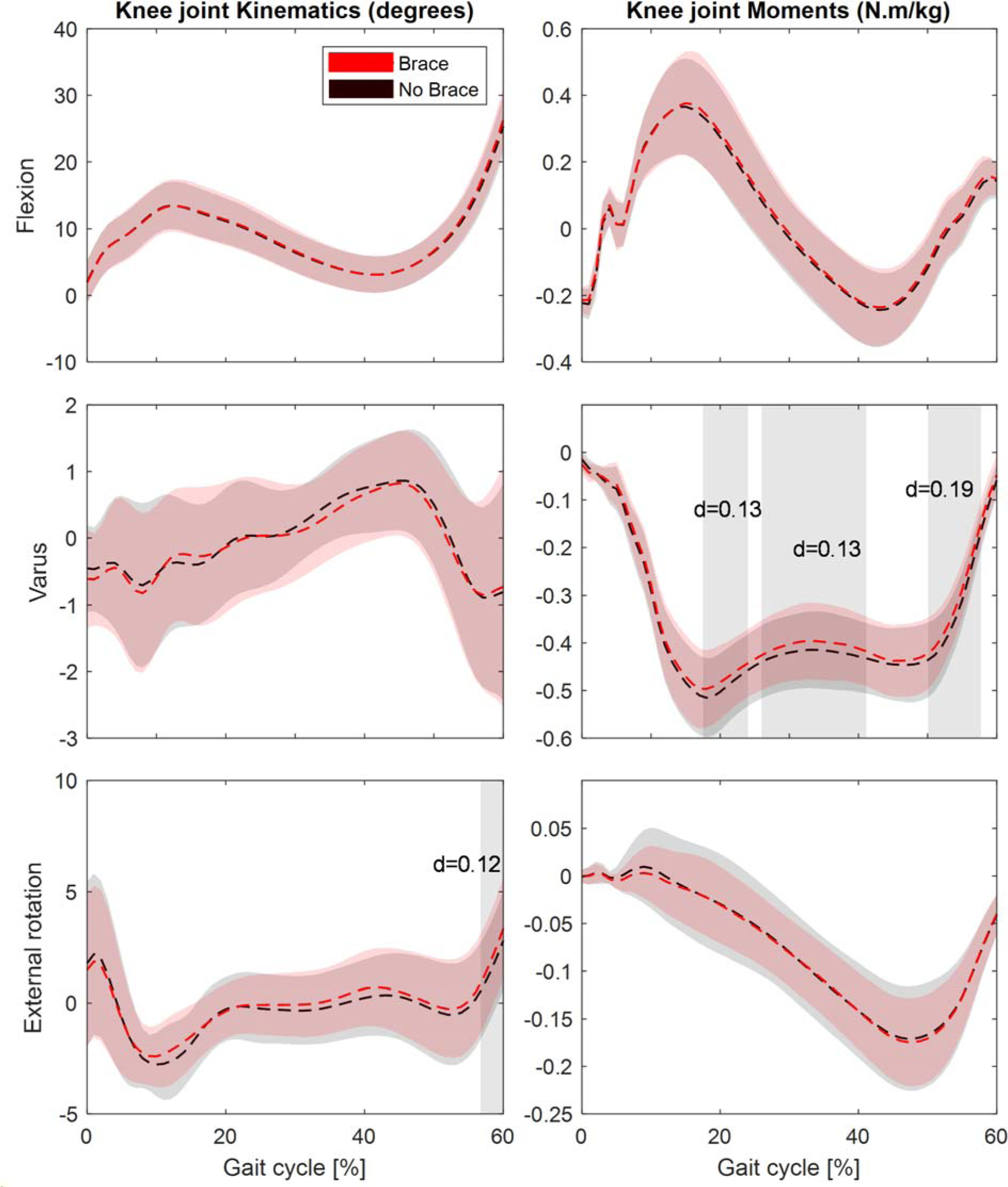
Knee joint angles and moments during the stance phase on the ACLR leg during walking. Brace condition is shown in red, and no brace in black. The shading represents the standard deviations. The grey bands represent periods of significant difference between the conditions with their mean Cohen d.

**Appendix 2:**
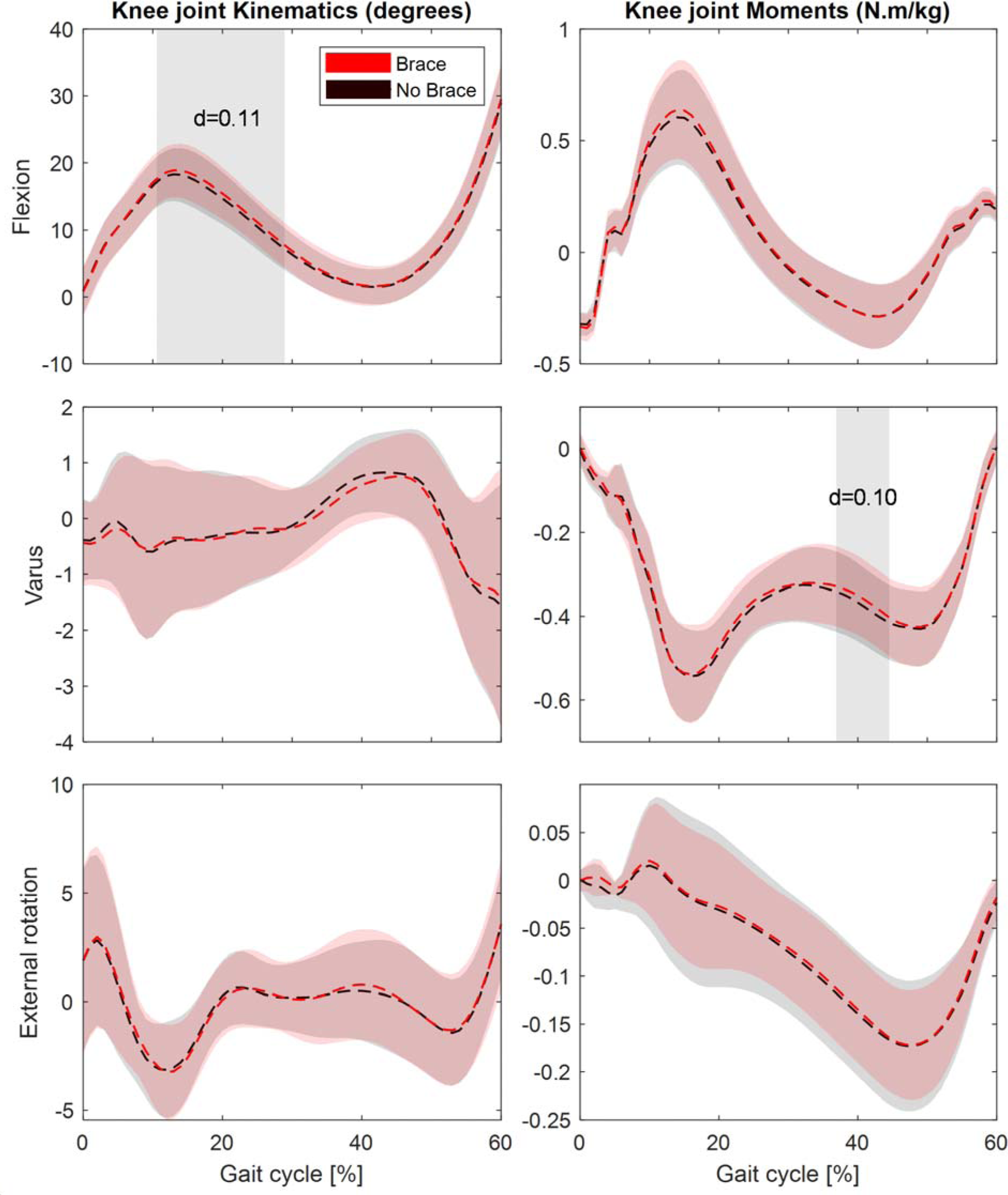
Knee joint angles and moments during the stance phase on the ACLR leg during fast walking. Brace condition is shown in red, and no brace in black. The shading represents the standard deviations. The grey bands represent periods of significant difference between the conditions with their mean Cohen d.

**Appendix 3:**
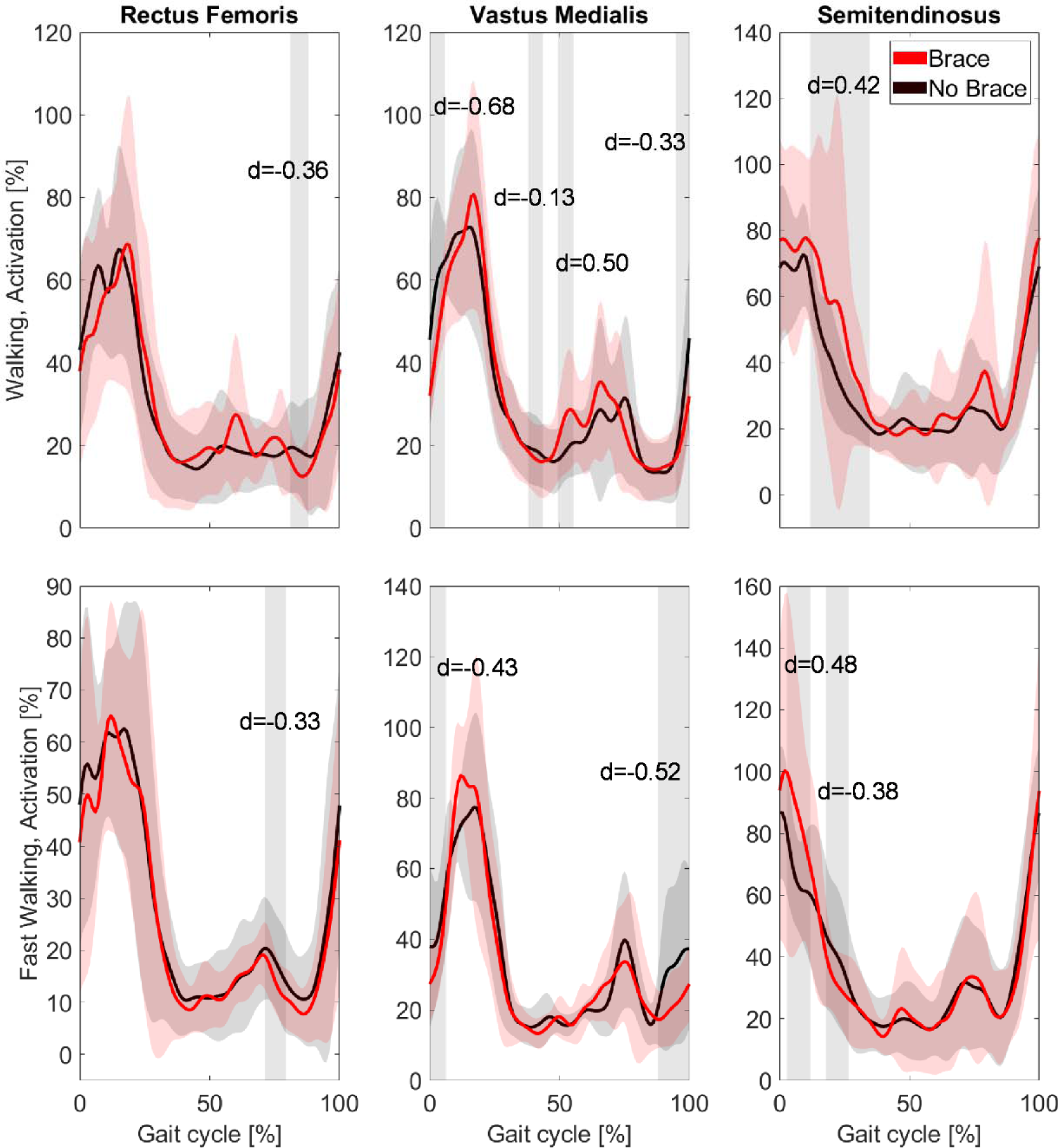
EMG activities during walking and Fast Walking task. Brace condition is shown in red, and no brace in black. The shading represents the standard deviations. The grey bands represent periods of significant difference between the conditions with their mean Cohen d. EMG is normalized with respect to peak value of the contralateral limb.

